# Mapping cellular targets of covalent cancer drugs in the entire mammalian body

**DOI:** 10.1101/2025.03.28.645991

**Authors:** Zhengyuan Pang, Verina H. Leung, Cailynn C. Wang, Ahmadreza Attarpour, Anthony Rinaldi, Hanbing Shen, Maria Dolores Moya-Garzon, Logan H. Sigua, Claire Rammel, Alexandra Selke, Christopher Glynn, Melaina Yender, Senhan Xu, Javid J. Moslehi, Peng Wu, Jonathan Z. Long, Maged Goubran, Benjamin F. Cravatt, Li Ye

## Abstract

As our understanding of biological systems reaches single-cell and high spatial resolutions, it becomes imperative that pharmacological approaches match this precision to understand drug actions. This need is particularly urgent for the targeted covalent inhibitors that are currently re-entering the stage for cancer treatments. By leveraging the unique kinetics of click reactions, we developed volumetric clearing-assisted tissue click chemistry (vCATCH) to enable deep and homogeneous click labeling across the 3D mammalian body. With simple and passive incubation steps, vCATCH offers cellular resolution drug imaging in the entire adult mouse. We combined vCATCH with HYBRiD imaging and virtual reality to visualize and quantify *in vivo* targets of two clinical cancer drugs, afatinib and ibrutinib, which recapitulated their known pharmacological distribution and revealed previously unreported tissue and cell type engagement potentially linked to off-target effects. vCATCH provides a body-wide, unbiased platform to map covalent drug engagements at unprecedented scale and precision.

## INTRODUCTION

The primary binding sites for drug molecules are critical determinants of drug efficacy and toxicity. Although molecular target identification strategies, such as chemoproteomics^1–5^, chemogenomic^6–9^, and thermal shift profiling^10–13^, have revolutionized our understanding of drug activities, mapping *in vivo* drug targets with spatial and cellular resolution remains challenging. This shortfall is even more pressing in light of the innovations in genomics and imaging techniques over the past decade that have enabled the characterization of tissue architecture at the single-cell level with spatial resolution^14–20^. This has led to a widening gap between our understanding of native biological systems and our ability to track *in vivo* drug actions within these systems, as the latter still primarily rely on low-resolution methods such as radioactive imaging. It is imperative that we develop methods to visualize *in situ* drug engagement with cellular resolution across tissue compartments and organ systems to better understand *in vivo* drug actions.

One promising avenue for imaging drug targets is fluorescence imaging, which has been widely used for visualizing endogenous biomolecules like DNA, mRNA, and proteins. However, directly imaging drug molecules conjugated with a fluorescent tag is often not suitable as a larger tag could distort drug activity and distribution *in vivo*. As an alternative approach, we recently developed a method to visualize drug molecules in tissue sections by combining tissue clearing with *in situ* click chemistry (CATCH)^21^. However, efforts to extend CATCH from thin sections to large, heterogeneous 3D tissue environments face significant challenges similar to those typically encountered by whole-mount immunostaining or *in situ* hybridization methods, such as the accumulation of probes at the tissue surface, uneven staining across the tissue depth, and insufficient labeling at the center of large samples^22–26^.

Here, we develop new CATCH capacities to enable rapid and homogenous *in situ* click labeling in ultra-large 3D tissues at the scale of a whole mouse body. This new technology leverages the unique characteristics of click chemistry by introducing pre-reaction copper saturation (PRCS) and repeated iterations of reaction (RIR). The resulting volumetric CATCH (vCATCH) method, when combined with HYBRiD (hydrogel-based reinforcement of three-dimensional imaging solvent-cleared organs (DISCO)) clearing^27^, is further highlighted by its marked simplicity as it only relies on easy and passive incubation steps to achieve whole-body labeling and imaging.

## RESULTS

### Establishing the principles for deep tissue click reactions

To achieve 3D click labeling, we turned to recent advances in whole-brain and whole-body immunostaining techniques^24,28–35^. As summarized by previous works^36–39^, two main strategies have been used to achieve deep tissue antibody penetration: (1) enhancing diffusion of the probe (e.g., antibody) into the tissue; and (2) minimizing probe-target interactions (e.g., antibody-antigen binding) during the diffusion stage while maximizing reaction efficiency at the staining stage, such that the large difference in reaction rates between the two stages will prevent probe depletion at the tissue surface and facilitate homogeneous tissue labeling^23,33,34^. Biophysics studies have demonstrated that changing the pH, salt concentration, and other physicochemical parameters can modulate antibody-antigen binding kinetics by 4-400 fold^40,41^. We hypothesized that we could adopt a similar strategy to enable 3D click reactions. CATCH relies on copper-catalyzed alkyne azide cycloaddition (CuAAC) to label target-bound drug molecules with fluorescent signals. As its name indicates, the kinetics of CuAAC strictly depend on the catalytic Cu(I)-ligand complex, but for ease of handling, a Cu(II) compound, typically CuSO_4_, is used during the diffusion stage and then a reducing agent (e.g., sodium ascorbate) is added *in situ* to generate Cu(I) at the binding stage. Compared to uncatalyzed conditions, the generation of catalytic Cu(I) can lower the energy barrier by 14.9-18.4 kcal/mol, or an equivalent of 7-8 orders of magnitude increase favoring the cycloaddition that attaches the fluorescent probe to the drug *in situ*^42^. Thus, by temporarily preventing the generation of Cu(I) from Cu(II) through withholding the reducing agent sodium ascorbate, in principle, we could modulate the probe-target reaction by 10^7–8^ fold between the diffusion and binding stages, which could improve the penetration depth of CATCH in large tissues.

For method development, we used the covalent monoamine oxidase (MAO) inhibitor pargyline, whose alkyne probe pargyline-yne has previously been characterized by activity-based protein profiling (ABPP) and 2D-CATCH studies (Figure S1A)^21,43^. Although CATCH has demonstrated homogeneous click labeling of N_3_-dye in 500 μm tissues^21^, when tested in mouse hemispheres, even with the aforementioned catalyst withholding strategy, only the surface of the mouse brain was labeled, a depth of 0.4 mm (Figure 1A). First, we used mouse brains treated with the fatty acid amide hydrolase (FAAH) inhibitor PF7845-yne (Figure S1A), another CATCH probe we previously established, to determine whether other CATCH components might have trouble diffusing through the tissue^21,44^. We removed individual reagents from the diffusion step and evaluated the resulting labeling depth. For instance, eliminating the N_3_-dye, the largest molecular weight component, in the diffusion step did not affect labeling depth, suggesting N_3_-dye diffusion is not a limiting factor (Figure S1B and S1C). By contrast, removing CuSO_4_ or ligand strongly reduced labeling depth in these conditions, suggesting that the catalyst is still likely the diffusion bottleneck (Figure S1B and S1C).

**Figure 1.**
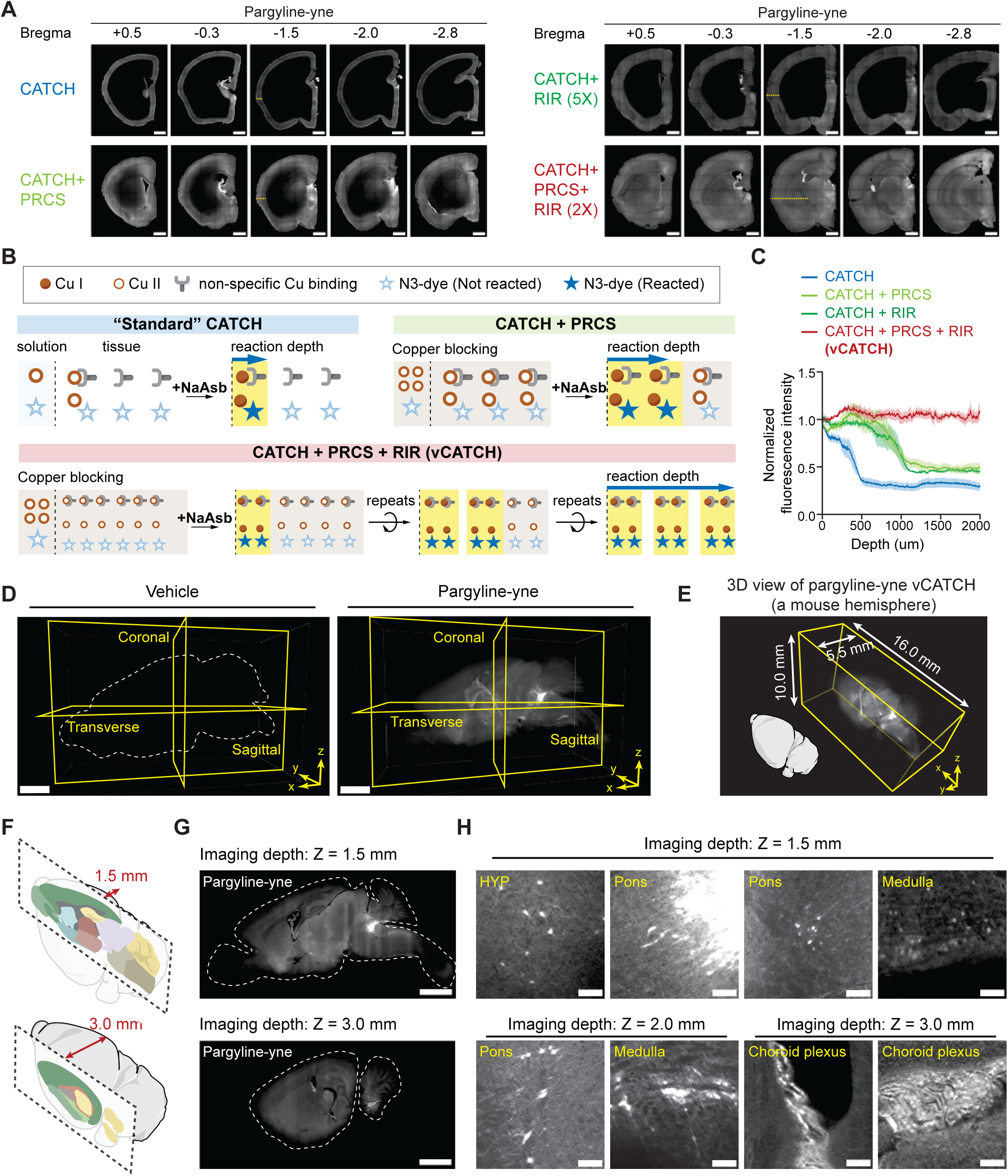
Establishing the principles for deep tissue click reactions. (A) Brain-wide characterization of click labeling in pargyline-yne-treated hemispheres (10 mg/kg, 1 h, i.p.). Pre-reaction copper saturation (PRCS) and repeated iterations of reactions (RIR, 1 hour per iteration) enhanced labeling depth and generated homogeneous click labeling across the whole hemisphere. (B) Schematics of copper competition in tissue. Copper binding sites could chelate copper and limit the available free copper for CuAAC catalyst conversion, thereby impairing click labeling efficiency deep in tissues. (C) Quantification of click labeling depth along the marked lines in Figure 1A. Four hemispheres in each condition. (D) Representative 3D rendering of vehicle and pargyline-yne treated hemispheres. (E) 3D rendering of pargyline-yne treated hemispheres showing the total imaging volume. (F) Schematics of imaging depth shown in Figure 1G, highlighting changes in different brain regions covered. (G) Sagittal 2D views of pargyline-yne treated hemispheres at imaging depth Z=1.5 mm and 3.0 mm as indicated in Figure 1F. (H) Zoomed-in views showing pargyline-yne enriched neurons and brain regions across the hemisphere. Data are plotted as mean±SEM. Scale bars: 1000 μm (A); 2000 μm (D and G); 100 μm (H)

Cu is well-known to bind multiple amino acid side chains (i.e., cysteine and histidine)^45–48^. We therefore hypothesized that the ubiquitous copper binding sites could consume free Cu(II) as it diffuses into the sample, resulting in insufficient catalyst conversion in deeper tissues (Figure 1B). Unlike typical staining in which increasing dye or antibody concentrations tends to result in nonspecific labeling and high background, high Cu(II) alone, as a catalytic precursor, should not lead to nonspecific labeling. Based on this theoretical prediction, we reasoned that excessive Cu(II) could be used to pre-saturate potential copper binding sites (therefore “blocking” them) to boost catalytic activity in deeper tissues. Importantly, this strategy should not increase nonspecific labeling or lead to high background. Indeed, we observed that incorporating a 10x Cu excess pre-reaction copper saturation (PRCS) step increased the reaction depth to 1 mm while maintaining the signal-to-noise ratio after a single one-hour reaction (Figure 1A-1C).

We next sought to modify CATCH to reach targets in tissues deeper than 1 mm. Although increasing reaction time could potentially increase labeling depth, prolonged reactions would deplete the reducing agent in the buffer, which could eventually lead to reactive oxygen species (ROS) build-up and side reactions^49–51^. Instead, we refreshed the click reaction cocktail every hour, a procedure we termed repeated iterations of reactions (RIR). In practice, PRCS plus two rounds of RIR resulted in full-depth, homogenous CATCH labeling across the mouse hemisphere (Figure 1A-1C, Supplementary Video 1). We term this approach volumetric CATCH (vCATCH).

### vCATCH quantitatively profiles brain-wide targets of Pargyline

After achieving brain-wide labeling, we employed vCATCH to unbiasedly map drug targets of pargyline. After intraperitoneal (i.p.) injection of pargyline-yne (10 mg/kg) or vehicle (non-drug treated) in wild-type mice, mouse hemispheres were harvested and cleared by HYBRiD. Hemispheres then underwent one day of PRCS and two rounds of RIR before imaging with lightsheet fluorescence microscopy (Figure 1D). Importantly, vCATCH allowed the use of high-throughput, paralleled passive processing to generate cohort-scale pargyline-labeled brains with marked reproducibility, critical for performing biologically meaningful analysis of its cellular targets across the whole brain (Figure 1E-1H, S1D, and S1E).

First, we manually verified pargyline probe binding in the brain based on previously reported regional targets^21^. These regions included distinct cellular structures in the pons and hypothalamus (Figure 1H and S1E). In addition, we found pargyline-yne-enriched cells in the choroid plexus (in the ventricle) and the medulla (Figure 1H and S1E). We also observed additional neuron-like structures in the midbrain (Figure S1E). These findings demonstrated that vCATCH maintained high fidelity in large tissue volumes and across the whole brain.

The throughput and reproducibility of vCATCH across cohort-scale samples also allowed automatic and unbiased quantification of drug targets beyond visual inspection. We applied a recently developed AI-based Cartography of Ensembles (ACE) pipeline to analyze these datasets (Figure 2A-2C, Supplementary Table 1)^52,53^. After registering the 3D volumes onto the Allen Brain Atlas (ABA)^54^ (Figure S2A), we first quantified the fluorescence intensity (indicating drug binding abundance) across all brain regions (Figure 2B). Intensity quantifications revealed high drug abundance in the pons, thalamus, and midbrain (summarized in Figure 2D and S2B), for example, in the locus coeruleus (LC) and nucleus accumbens (ACB) (Figure 2D).

**Figure 2.**
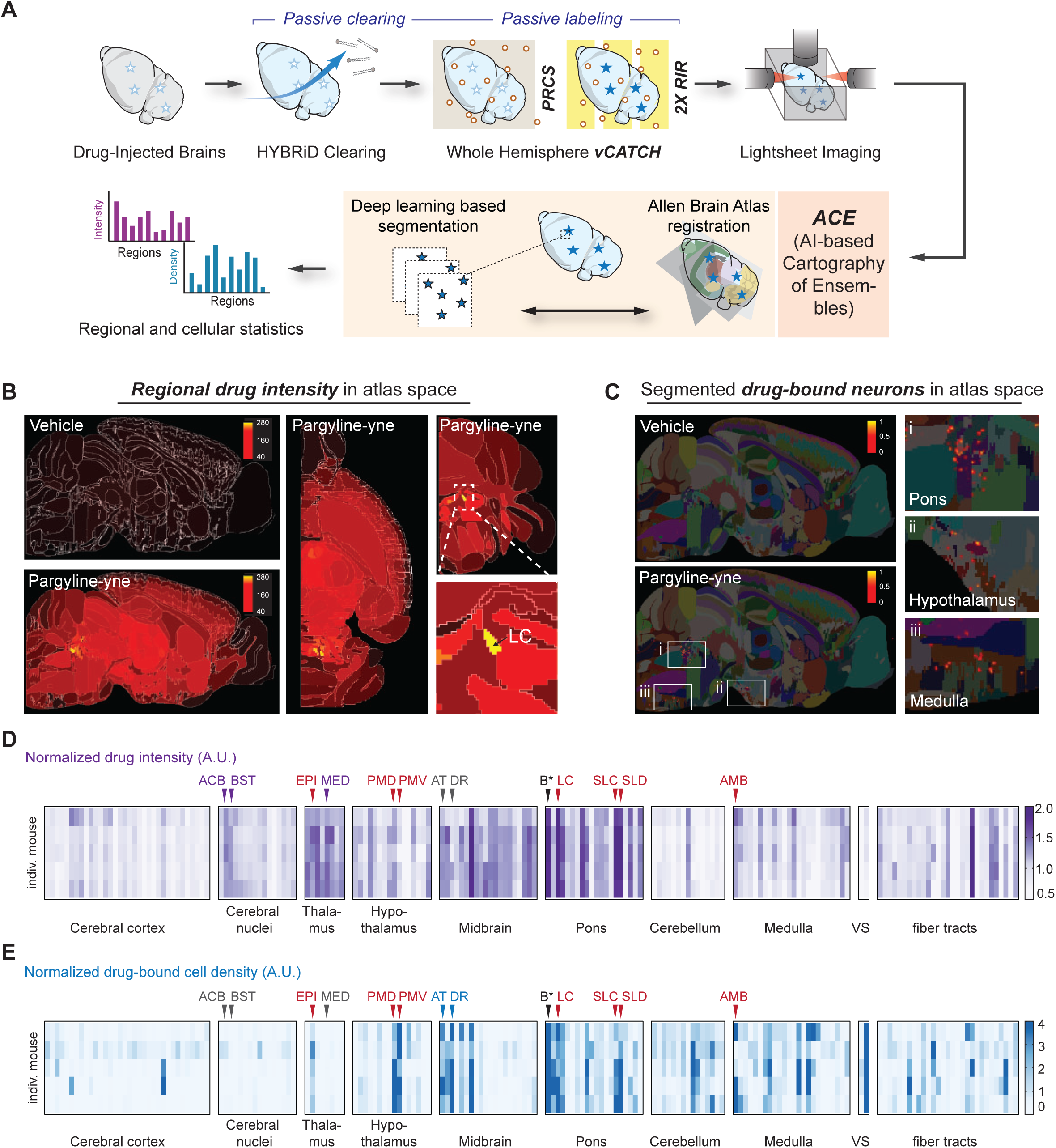
ACE analysis of pargyline-yne targets across the hemisphere. (A) Schematics of the ACE analysis pipeline for brain wide pargyline-yne (10 mg/kg, 1 h, i.p.) target identification. (B) Representative regional drug intensity quantification in atlas space. White boundary indicates brain regions by Allen Brain Atlas (ABA). The LC showed the highest overall labeling intensity in pargyline-yne hemispheres. Raw intensity values indicated by scale bar. (C) Representative segmented neurons in atlas space. Known pargyline-yne-bound neurons in the pons and hypothalamus were used as positive controls. Additional neuronal structures (i.e. in the medulla) were identified. The scale bar indicates raw segmented neuron density in each 10x10x10 mm voxel. (D-E) Heatmap of normalized pargyline-yne labeling intensity (D) and normalized pargyline-yne positive neuron density (E). A total of 5 pargyline-yne-treated mice (rows) and 184 brain regions (columns) were included for analysis. For individual mice, intensity and density values were normalized by the average intensity and density of all the regions from each mouse. Purple labels indicate regions with only elevated intensity. Blue labels indicate regions with only elevated neuronal density. Red labels indicate regions with shared elevated intensity and neuronal density. Barrington’s nucleus (B*) was later found to be a false positive (See Figure S2F). The full dataset and acronyms can be found in Supplementary Table 1. EPI, epithalamus; PMD, dorsal premammillary nucleus; PMV, ventral premammillary nucleus; AT, anterior tegmental nucleus; SLC, subceruleus nucleus; SLD, sublaterodorsal nucleus; AMB, nucleus ambiguus; VS, ventricular system.

Leveraging the cellular resolution imaging provided by vCATCH, we next sought to unbiasedly quantify the cell targets of pargyline. Empowered by a large training dataset and AI-based segmentation algorithms, the ACE pipeline was used to identify brain-wide pargyline-yne-positive cells (Figure 2C, S2C, and S2D). After segmentation, the cell density patterns generally agreed with fluorescence intensity profiles. Since pargyline targets both MAO-A and MAO-B with similar affinities^43^, we performed MAO-A and MAO-B immunofluorescence and confirmed that pargyline-yne-targeted cells overlap with either or both MAO-A/B protein expression, consistent with the known affinity of pargyline on these two targets (Figure S2E)^43^.

Interestingly, unbiased ACE analysis also revealed unexpected differences between fluorescence intensity and cell counts. For example, the ACB, bed nuclei of the stria terminals (BST), and the medial group of the dorsal thalamus (MED), showed extensive drug intensity, but almost no neuronal structures were detected in these areas, indicating drug binding may be associated with non-soma structures (Figure S1E, 2D, and 2E). Conversely, MAO-B positive drug-bound cells were identified in the dorsal raphe (DR) and the anterior tegmental nucleus (AT) of the pons (Figure S1E, 2D, 2E, and S2E) despite the lack of overall fluorescence intensity increase, suggesting sparse neuronal populations were targeted by pargyline in these regions.

Taken together, by using a pargyline probe as a proof-of-concept, we demonstrated that vCATCH is compatible with clearing methods and high-throughput computational pipelines to unbiasedly quantify regional enrichment and cellular targets of covalent drugs in the mouse brain, which was not possible with traditional methods.

### vCATCH profiles whole-body targets of covalent cancer drugs

We next sought to expand vCATCH target mapping to whole animals across all organ systems. We selected two widely used cancer drugs for this study: afatinib (Gilotrif^®^) and ibrutinib (Imbruvica^®^), both covalent targeted kinase inhibitors (TKIs)^55,56^. Since their landmark approval in 2013, the epidermal growth factor receptor (EGFR) inhibitor afatinib and Bruton’s tyrosine kinase (BTK) inhibitor ibrutinib, respectively, have transformed the clinical landscape of non-small cell lung cancer (NSCLC)^57,58^ and B cell malignancies^59–61^. Despite their popularity, however, there are several well-documented side effects of TKIs that are poorly understood. For example, ibrutinib is associated with cardiovascular and bleeding risks^62,63^. Although many studies have been conducted to understand the protein targets (or off-targets) of ibrutinib^64–68^, the *in vivo* tissue(s) or cell types responsible for such toxicities remain elusive. The ability to unbiasedly profile TKI targets *in vivo* at cellular resolution could provide crucial insight to bridge this gap.

We injected previously established clickable probes of ibrutinib-yne (10 mg/kg, i.p.) and afatinib-yne (10 mg/kg, i.p.) in wild-type mice (Figure S3A)^69^. These probes largely retain the pharmacokinetics and tissue distribution of the parent compounds (Figure S3B and S3C). We established a vCATCH protocol with 4-day PRCS and 5 rounds of RIR to homogenously label a whole HYBRiD-cleared, one-week-old mouse with a total volume of more than 38x18x12mm (Figure 3A-3D). Labeling specificity was again confirmed by controlling for each vCATCH component in the lung or the spleen (canonical EGFR+ or BTK+ tissue, respectively, Figure S3D-S3G).

**Figure 3.**
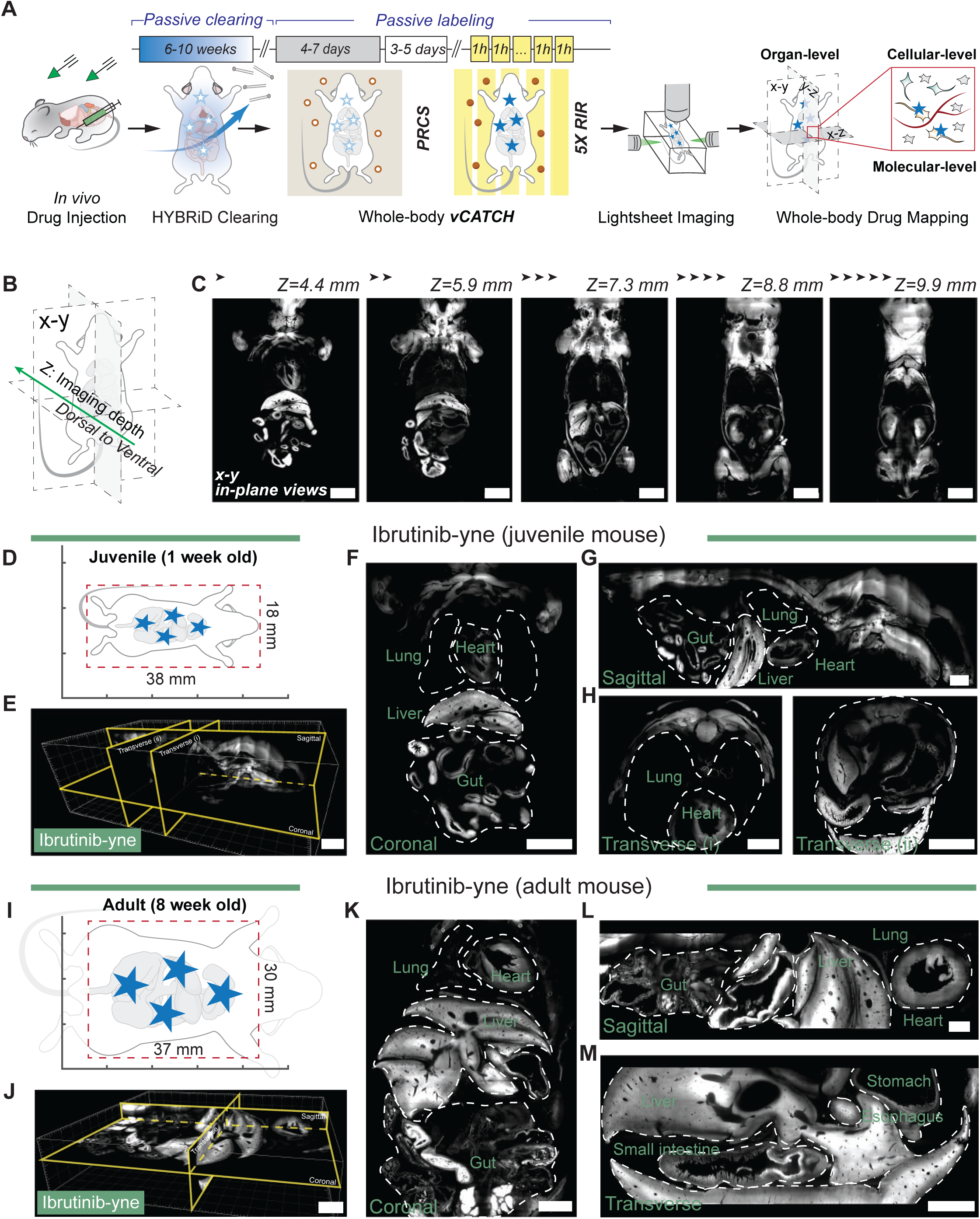
vCATCH reveals whole-body TKI distribution with high spatial resolution. (A) Workflow of vCATCH whole-body labeling with estimated time of each step. (B) Schematic of imaging depth. (C) Coronal (x-y) views of vCATCH whole-body labeling in ibrutinib-yne (10 mg/kg, 1 h, i.p.) treated juvenile mouse across different z-depths. (D) Schematic of lightsheet acquisition FOV, represented by red dotted box, for juvenile whole body. (E) Representative ibrutinib-yne distribution overviews in a juvenile mouse with digital plane slicing in 3D volume. (F-H) Coronal (F), sagittal (G), and transverse (H) views of a juvenile mouse as shown in Figure 3E. (I) Schematic of lightsheet acquisition FOV, represented by a red dotted box, for adult mouse torso. (J) Representative ibrutinib-yne distribution overview in an adult mouse torso with digital plane slicing in 3D volume image. (K-M) Coronal (K), sagittal (L), and transverse (M) views of adult mouse as shown in Figure 3J. Dashed lines indicate body boundary. Scale bars: 4000 μm (C, E, F, J, K); 2000 μm (G, H, L, M).

We then performed whole-body lightsheet imaging of these 1-week-old mouse bodies (Figure 3B-3H, S3H, S3J-S3M) and visualized them through virtual reality (VR) (Supplementary Video 2 and 3)^70^. The performance of vCATCH in young mice motivated us to expand its capacity to adult bodies. Enhanced by 7-day PRCS and 8 rounds of RIR (Figure 3A), we labeled ibrutinib-yne in an adult mouse with a total volume of 70x30x20 mm. Although this whole volume exceeded the field of view (FOV) of most commercial lightsheet systems, we were able to focus on the center of the torso (37x30x8 mm, Figure 3I) and examined drug distribution of major organs (Figure 3J-3M, and S3I).

Globally, the binding of both TKIs was highly abundant in the liver and the gastrointestinal tract (Figure 3F, 3K, S3K and S4). In the upper chest, ibrutinib showed elevated heart binding compared to the lung, whereas the opposite enrichment was observed with afatinib (Figure 4A, 4F and 4K). These findings are consistent with the general distributions of these two drugs based on established autoradiography data from the Food and Drug Administration^71,72^.

**Figure 4.**
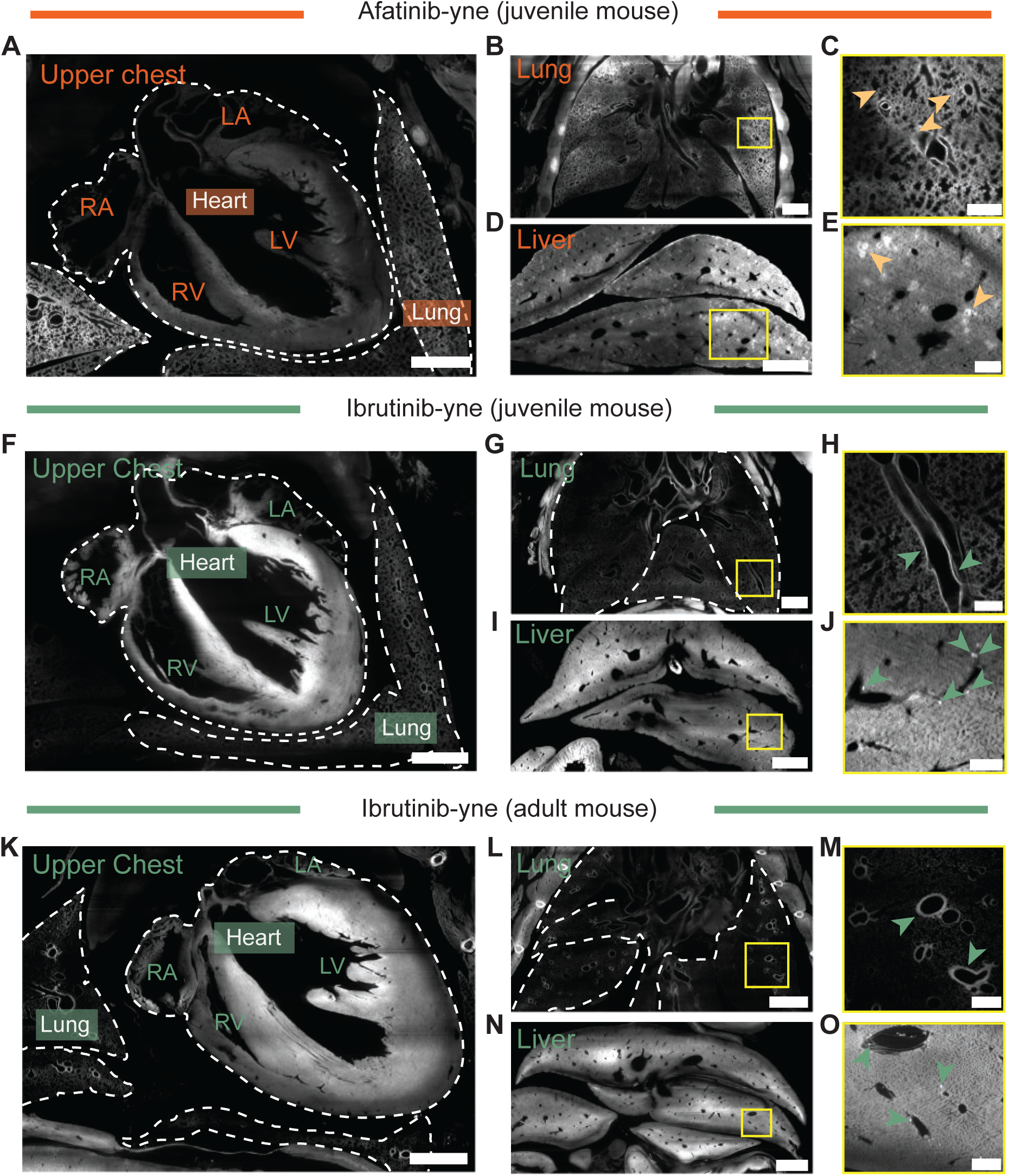
vCATCH reveals intra-organ TKI distribution. (A-E) Afatinib-yne binding in the upper chest (A), lung (B and C), and liver (D and E) of juvenile mice. (F-J) Ibrutinib-yne binding in the upper chest (F), lung (G and H), and liver (I and J) of juvenile mice. (K-O) Ibrutinib-yne binding in adult mice. Figures showing the upper chest (K), lung (L and M), and liver (N and O). Arrows indicate drug-enriched structures. Dashed lines indicate tissue boundary. Scale bars: 1000 μm (A, B, F, G); 500 μm (C, D, H, I, M, O); 100 μm (E, J); 2000 μm (K, L, N).

The resolution of vCATCH further allowed us to examine intra-organ drug distribution in detail. In several organs, afatinib and ibrutinib displayed similar intra-organ distributions; for example, both drugs were selectively enriched in the villus in the small intestine (Figure S4). Higher-magnification confocal imaging further revealed detailed drug-bound structures across organs (Figure 5A-5F). For example, both drugs strongly bound to circular structures in the bone (Figure 5A and 5D), which were also observed in the spleen (Figure S4). Subsequent lectin staining identified these structures as vasculatures (Figure S5A and S5B). Considering that BTK expression is generally absent in blood vessels^73^, we suspect the vasculature associated binding may be implicated in the off-target toxicity associated with ibrutinib^62,63^.

**Figure 5.**
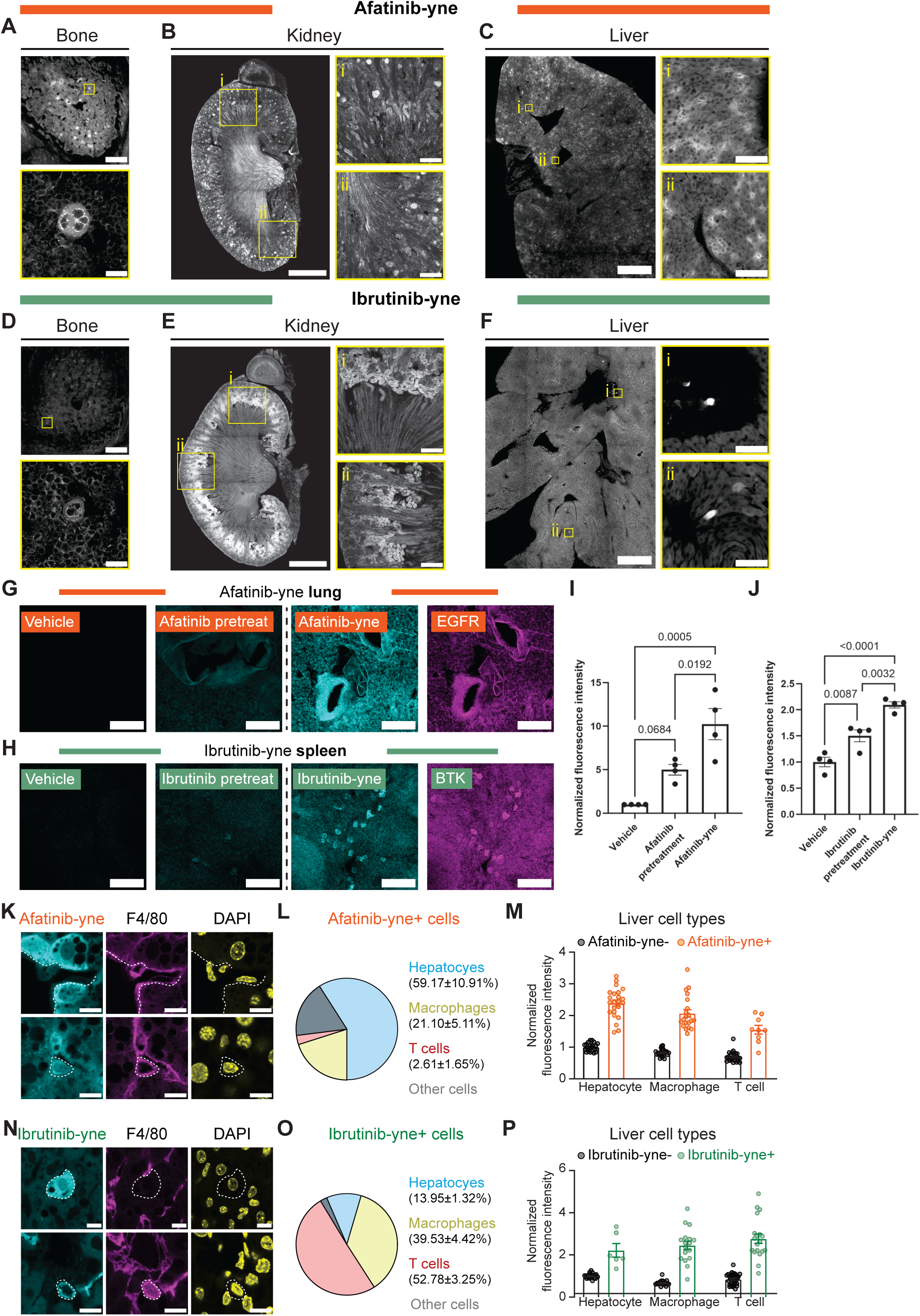
Cell type characterization of TKI engagement in drug-enriched tissues. (A-F) Afatinib-yne (A-C) and ibrutinib-yne (D-F) characterizations in the bone (A, D), kidney (B, E), and liver (C, F). (G) Afatinib pretreatment (25 mg/kg, 1 h, i.p.) in the lung co-stained with EGFR. (H) Ibrutinib pretreatment (25 mg/kg, 1 h, i.p.) in the spleen co-stained with BTK. (I-J) Quantification of afatinib-yne labeling intensity of EGFR-positive pixels in the lung (I) and ibrutinib labeling intensity of BTK-positive pixels in the spleen (J). n = 4 mice for each condition. One-way ANOVA with Tukey’s multiple comparisons test. p-values plotted in the graph. (K) Representative images of afatinib-yne+ hepatocytes (top row) and afatinib-yne+ macrophages (F4/80+, bottom row). (L-M) Afatinib-yne positive cell type (L) and cell type specific intensity (M) quantifications. n=4 biological replicates, 4-6 FOV were acquired for each mouse. (N) Representative images of ibrutinib-yne+ hepatocytes (top row) and ibrutinib-yne+ macrophages (F4/80+, bottom row). (O-P) Ibrutinib-yne positive cell type (O) and cell type specific intensity (P) quantifications. n=4 biological replicates, 4-6 FOV were acquired for each mouse. All values are shown as mean ± SEM. Dashed lines represent the cell boundary. Detailed cell type quantification is in Supplementary Table 1. Scale bars: 200 μm (A, C, global view; B, E, zoomed-in view; G, H); 20 μm (A, C, zoomed-in view); 1000 μm (B, E, global view); 500 μm (C, F, global view); 50 μm (C, F, zoomed-in view); 10 μm (K, N).

In other organs, we observed distinct target distributions between these two drugs. For example, robust tissue enrichments in the inner stomach lining and along the nasal tract were associated with ibrutinib but not afatinib (Figure S4A and S4B). Furthermore, we discovered widely spread afatinib targets in the lung, whereas pulmonary ibrutinib labeling was restricted along the bronchi (Figure 4B, 4C, 4G, 4H, 4L, and 4M).

Interestingly, although both drugs were bound to the kidney, they displayed different intra-organ patterns (Figure S4). Afatinib was associated with glomeruli in the cortex and near the cortex-medulla interface, whereas ibrutinib was more localized to cortical tubular-like structures (Figure 5B and 5E). Similarly, our global organ-level imaging (Figure 4D, 4I, and 4N) showed that both TKIs were enriched in the liver, in agreement with the literature^71,72^. However, higher resolution vCATCH revealed differences in their intra-organ distribution: afatinib showed crown-like signals throughout the hepatic tissue, whereas ibrutinib was associated with sparsely scattered individual cells, as demonstrated by both lightsheet (Figure 4E, 4J, and 4O) and confocal imaging (Figure 5C and 5F).

### Multiplexed molecular characterizations of drug-enriched cellular structures

Guided by the drug distribution patterns, we next sought to molecularly characterize drug-targeted cells. We first focused on afatinib binding in the lung and ibrutinib binding in the spleen based on their canonical “on-target” engagements^73^. As expected^69^, afatinib-yne and ibrutinib-yne signals were primarily on EGFR+ and BTK+ cells, respectively, as determined by immunostaining (Figure 5G and 5H). Importantly, pretreating the animals with the parent compound of each probe robustly reduced probe labeling (Figure 5G-5J), further confirming the specificity of these probes.

In the liver, we observed enriched yet distinct binding patterns of afatinib-yne and ibrutinib-yne (Figure 5C and 5F). We next performed molecular analysis to determine the targeted cell types of each probe. First, consistent with high EGFR expression in the hepatocytes^73^, we found that the majority (59.17±10.91%) of afatinib-bound cells were hepatocytes, followed by small fractions of macrophages (F4/80+) and T cells (CD3e+) (Figure 5K-5M, and S5C). By contrast, only a small fraction (13.95±1.32%) of ibrutinib-bound cells were hepatocytes, while a larger proportion (39.53±4.42%) of these cells were F4/80+ macrophages (Figure 5N-5P), consistent with known BTK expression in liver macrophages^73^. Interestingly, the largest fraction (52.78±3.25%) of ibrutinib-bound cells were CD3e+ T cells despite their low BTK expression (Figure 5O and S5C-E)^73^, suggesting non-BTK engagement could be involved (see Discussion).

Beyond the liver, we were also able to identify and quantify afatinib-enriched immune cell populations in other tissues such as the lung (Figure S5F-S5H). In summary, our vCATCH data revealed that the two TKIs are enriched in distinct cell types within individual tissues and organs, highlighting vCATCH’s capacity to profile drug targets across global, organ, tissue, and cellular scales.

## DISCUSSION

Covalent TKIs have reentered the stage for cancer treatment and broadened the horizon for the druggable space as demonstrated by the recent developments of mutant selective EGFR inhibitor osimertinib, and KRAS G12C inhibitors sotorasib and adagarasib^55^. However, despite the reignited enthusiasm, the concern for toxicity (for example, cardiotoxicity and bleeding for ibrutinib) calls for continuous innovations in characterizing their *in vivo* targets to guide future covalent drug development.

Here, by treating samples with excessive copper (PRCS) and performing multiple rounds of click reactions (RIR), we established a new method, vCATCH, which enabled robust covalent TKI mapping across the whole mouse body. This method only requires simple passive incubations and therefore offers reliability, reproducibility, and high throughput for cohort-scale biological and pharmacological analyses. Using existing pharmacokinetic data from the Food and Drug Administration (FDA) as a reference, vCATCH confirmed organ-level drug distribution. Critically, vCATCH enabled us to identify drug engagement with intra-organ cellular resolution that is typically inaccessible through conventional methods. The capacity to resolve drug engagement across molecularly defined cell populations, tissue architectures, and organ systems, once combined with single-cell and spatial omics tools, presents transformative opportunities for spatially defined, cell-type-specific drug action characterization.

Our work so far has focused on wild-type mice for method development. By examining drug binding in healthy mice, we uncovered patterns potentially associated with off-target engagement and toxicities. For example, ibrutinib is well known to be associated with cardiac toxicity such as atrial fibrillation (AF), but the mechanism behind this association remains incompletely understood^64–67^. Although animal studies suggested AF induction would likely require chronic administration^66^, our study revealed clear ibrutinib enrichment in the heart after a single injection, suggesting an intrinsic enrichment of ibrutinib for the cardiovascular system. It would be of great interest to further examine ibrutinib engagement in AF models to better understand its cardiotoxicity mechanism. In addition to AF, bleeding is another major ibrutinib risk factor^62,63^. Despite low BTK expression in endothelial cells^73^, we observed ibrutinib engagement in the vasculature, a finding that could have implications for ibrutinib-associated hypertension and help guide subsequent molecular identification work to further characterize unique ibrutinib targets in endothelial cells^75^.

Moreover, we observed ibrutinib binding in T cells that normally do not express BTK. This binding suggested non-BTK targets of ibrutinib at the dose we used in our studies. Indeed, previous ABPP studies in Ramos cells have identified human epidermal growth factor receptor 2 (HER2)^69^ as an ibrutinib target. *In vitro* kinase activity profiling also identified interleukin-2 inducible T-cell kinase (ITK) and Janus kinase 3 (JAK3) with nM affinity to ibrutinib^68^. All of these non-BTK affinities could contribute to our observed T cell binding.

On the technology side, vCATCH represents a new strategy to use biorthogonal click chemistry to label large tissues, in contrast to most mainstream labeling methods (e.g., immunostaining, DNA/RNA hybridization, etc) that primarily rely on biological affinities, such as antibody-antigen, biotin-streptavidin, and complementary DNA/RNA interactions. Click chemistry allows us to easily control labeling kinetics by simple buffer exchanges, achieving uniform probe penetration in ultra-large tissue without delicate procedures or apparatus. The covalent click chemistry linkage also means that the same sample can withstand harsh and prolonged processing and imaging operations, opening numerous possibilities to multiplex vCATCH with other spatial omics methods.

Finally, it is a major challenge in whole-mount tissue staining to balance the high probe concentration (such as antibody) required for tissue penetration and its associated background resulting from the nonspecific interactions between the probe and endogenous biomolecules. In this regard, the biorthogonal nature of click chemistry could better tolerate a wide range of labeling conditions without increasing noise. Currently, click chemistry-based imaging studies are primarily associated with examining transcription/translation^77,78^. By demonstrating vCATCH’s capacity to map covalent drug binding across the entire mouse body, we propose that similar principles can be applied in ultra-large 3D tissues to visualize substrates including but not limited to protein^79^, nucleic acid^80^, glycan^81^, lipid^82^, and drug molecules^21^. For example, a recent study (Click3D) has demonstrated that the original CATCH or similar strategies can label endogenous RNA and dyes in the mouse brain and tumor^83^, suggesting such approaches could be adopted in future biomedical studies, although it was limited to small, individual organs.

## LIMITATIONS OF THE STUDY

Focusing on method development, our current study used healthy wild-type mice, in which the engagement of TKIs could differ from tumor-bearing animals. Given the robustness and simplicity of the vCATCH pipeline, however, we believe other researchers with oncology expertise could readily apply vCATCH in various cancer models. We also note that due to the size limit of the current lightsheet microscopes, our imaging scope is limited to the center of adult mouse torsos. We foresee that the challenge can be addressed with emerging hardware and reconstruction pipelines^31,84,85^. Moreover, all clickable TKI probes used in this study have been rigorously validated by published medicinal chemistry and chemoproteomics studies^69^. However, we acknowledge that generating alkyne derivatives with similar absorption, distribution, metabolism, and excretion (ADME) properties of the original drugs requires dedicated efforts. We note that our ibrutinib-yne probe has a higher liver distribution than the parent drug, despite the latter being known to be enriched in the liver based on the FDA^72^. This difference could confound our interpretations of ibrutinib-bound immune cells (Figure 5). Lastly, as CATCH readout can potentially reflect both on-and off-target binding and metabolite intermediates (i.e., glutathione conjugation for cysteine-targeting drugs), it is important to note that the design of future CATCH probes for other drugs must be verified by similar PK and target identification approaches; although it is becoming common to generate and characterize clickable alkyne analogs during the development of new covalent inhibitors.

## RESOURCE AVAILABILITY

### Lead Contact

Further information and requests for resources and reagents should be directed to and will be fulfilled by the lead contact, Li Ye

### Materials availability

All unique/stable reagents generated in this study are available from the lead contact with a completed materials transfer agreement.

### Data and code availability

All data are available in the main text or the supplementary materials. Any additional information required to reanalyze the data reported in this paper is available from the lead contact upon request. Relevant code for the study is available from https://github.com/yelabscripps/vCATCH.

## ACKNOWLEDGMENTS

We thank all members of the Ye lab and the Dorris Neuroscience Center for their support and feedback. We thank Barry Sharpless for suggestions. This work was supported by NCI IMAT program (CA281918 to L.Y. and B.F.C), NIH Director’s New Innovator Award (DK128800 to L.Y.), and NIDA (DA059393 to L.Y.). L.Y. was also supported by the HHMI, NIDDK, NIMH/BRAIN, Chan Zuckerberg Initiative, and the Dana, Whitehall, Baxter, and Abide-Vividion Foundations. J.J.M received funding from the NIH (R01HL141466, R01HL155990, R01HL156021, P01HL141084). Z.P. and H.S. were supported by the Dorris Scholar Award. L.H.S is supported by an NIH Medical Scientist Training Grant T32GM007198-49. M.D.M.-G. was supported by Stanford Wu Tsai Human Performance Alliance. P.W. is supported by the NIH (R35GM139643).

## AUTHOR CONTRIBUTIONS

L.Y., Z.P. conceived the study. L.Y., Z.P., V.H.L., P.W., J.J.M., J.Z.L., M.G., and B.F.C. designed the experiments. Z.P., V.H.L., C.C.W., C.G., L.H.S., M.Y., A.S., and S.X. performed the experiments and analyzed the data. A.A., A.R., H.S., and M.G. performed the ACE analysis. Z.P., V.H.L., and M.D.M.-G. performed the PK study. Z.P., V.H.L., and C.R. performed the parental drug blocking study. L.Y. and B.F.C. acquired funding. L.Y. supervised the study. L.Y., Z.P., V.H.L., and C.C.W. wrote the manuscript with input from all authors. All authors reviewed and provided feedback on the manuscript.

---

**Figure S1.**
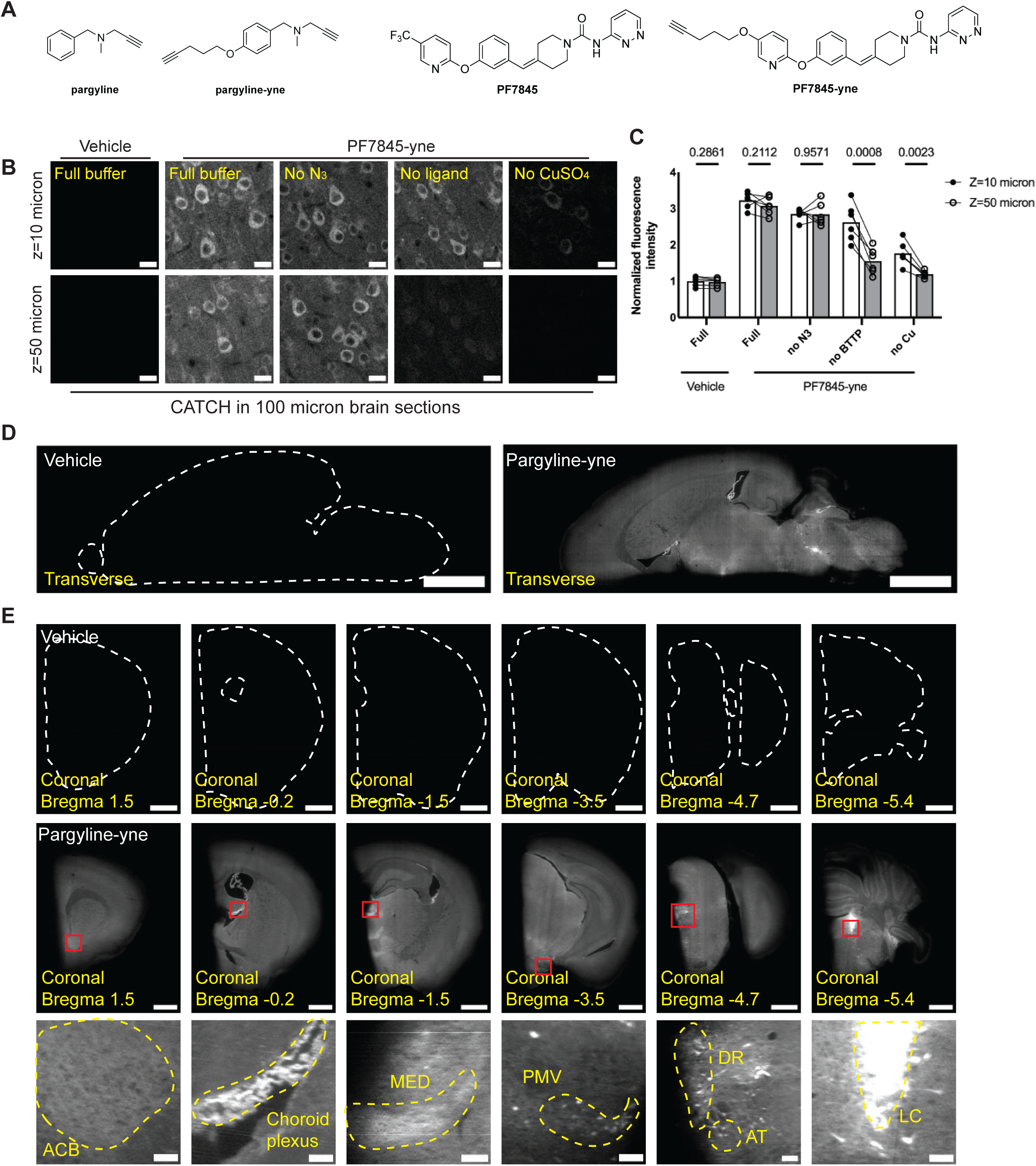
vCATCH development, related to Figure 1. (A) Chemical structure of the monoamine oxidase A/B (MAO A/B) inhibitor pargyline, fatty acid amide hydrolase (FAAH) inhibitor PF7845, and their alkyne analogs used in the study. Both analogs have been verified in previous CATCH and ABPP studies. (B) CATCH in 100-micron vehicle or PF7845-yne treated (1 mg/kg, 1 h, i.p.) brain sections. Individual components (Cu, ligand, N_3_) were removed during the click reaction incubation. Representative images were taken at the primary somatosensory cortex (S1, imaging depth Z=10 and 50 microns). (C) Quantification of CATCH intensity in Figure S1B. Six tissue sections (one field of view per section) were used for each condition. Intensity at Z=10 or 50 microns was measured and then normalized to the average vehicle labeling intensity at Z=10 or 50 microns, respectively. Paired t-test in each condition. P-values are plotted in the graph. n = 6 tissue sections for each condition. (D) Transverse 2D views of hemispheres in Figure 1E. (E) Coronal 2D views of hemispheres in Figure 1E. Zoomed-in views showing pargyline-yne enriched structures across the whole brain. Statistics determined by two-tailed paired t-test in (C). Dashed lines indicate tissue boundary. Scale bars: 20 μm (B); 2000 μm (D); 1000 μm (E, coronal view); 100 μm (E, zoom-in view) ACB, nucleus accumbens; MED, medial group of the dorsal thalamus; PMV, ventral premammillary nucleus; DR, dorsal nucleus raphe; AT, anterior tegmental nucleus; LC, locus coeruleus.

**Figure S2.**
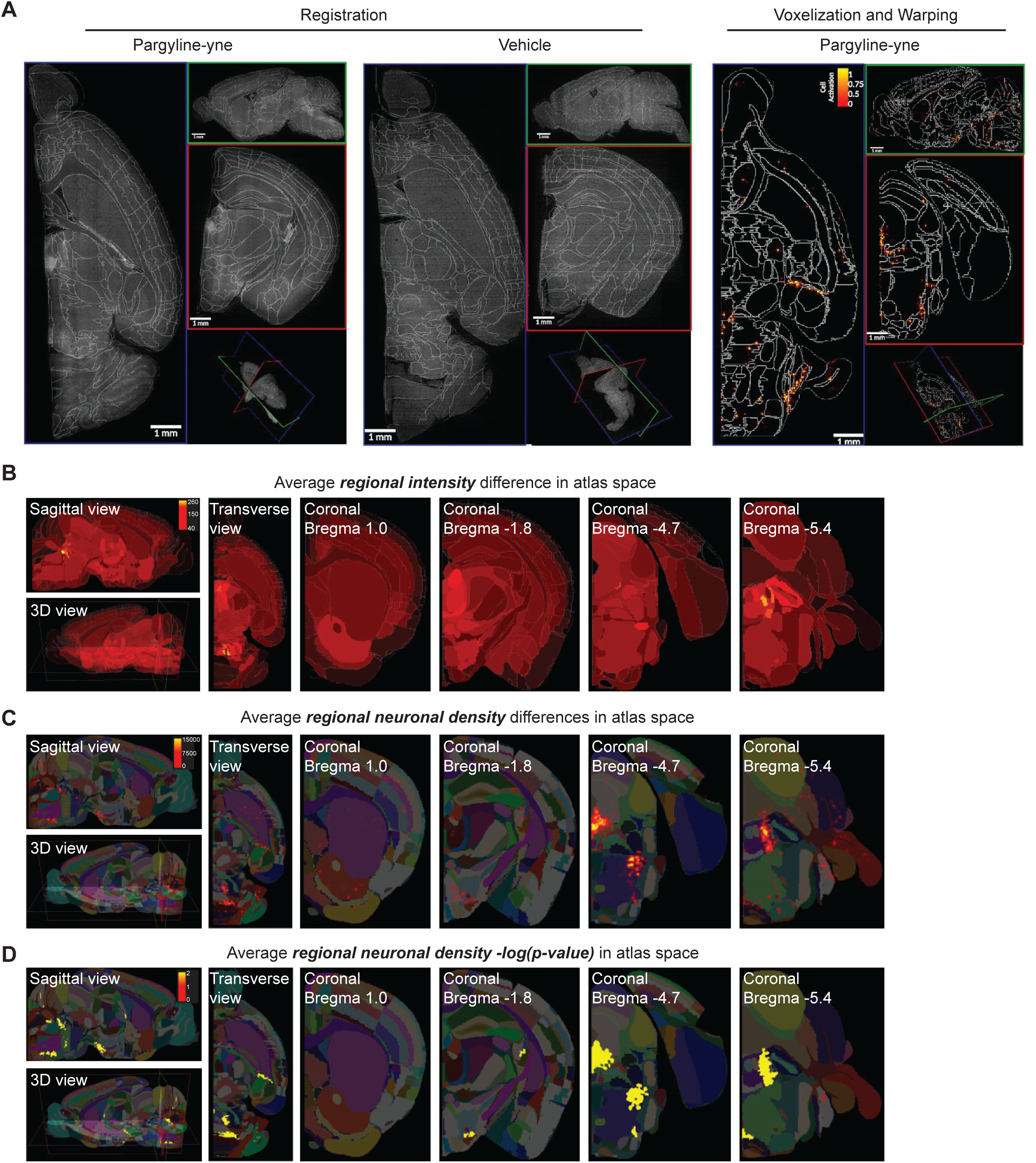
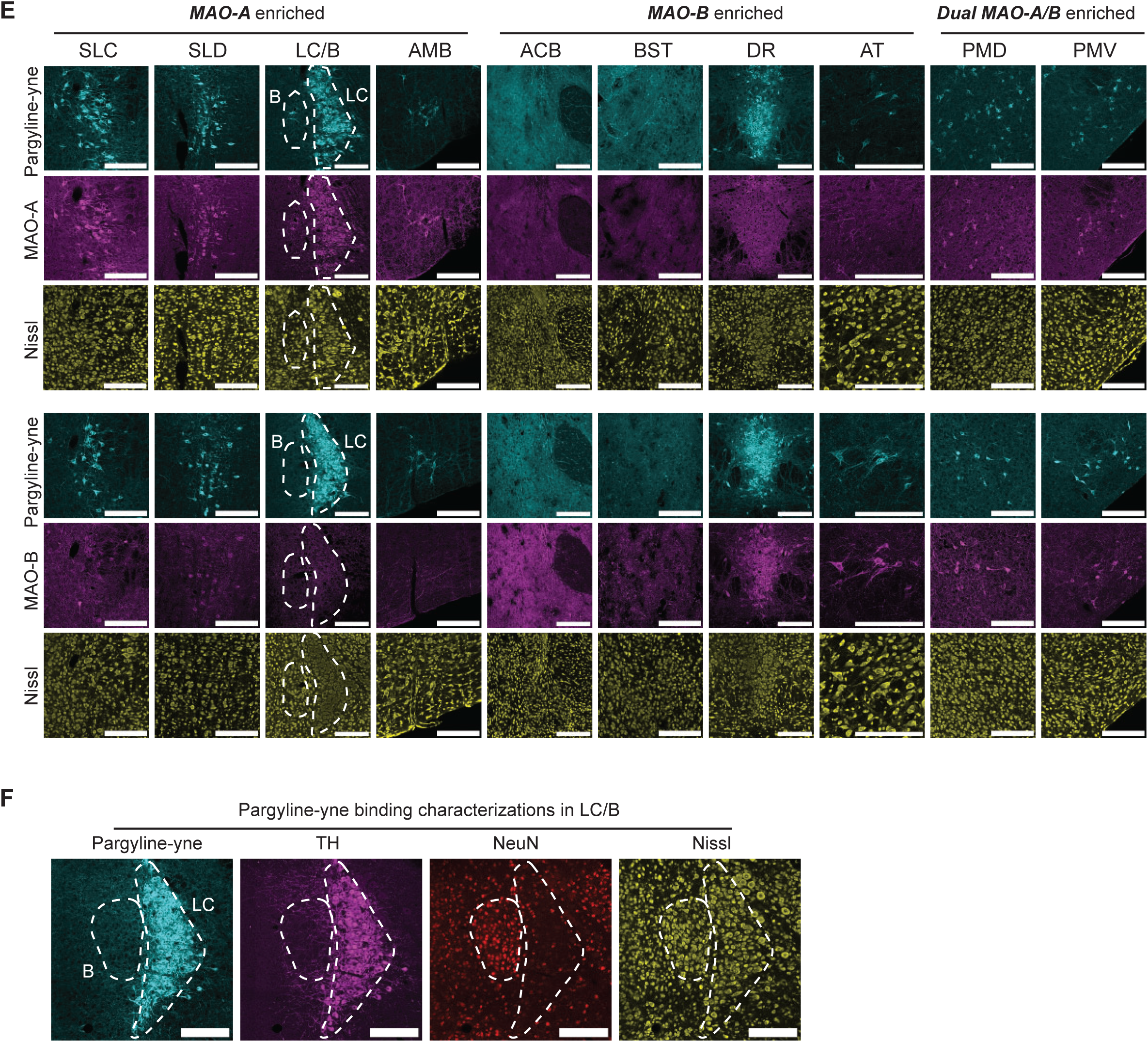
ACE analysis and validation, related to Figure 2. (A) Evaluation of ACE registration and warping algorithms. Lightsheet data were registered to the ABA using ACE. Left and right panels show data overlaid on labels for two subjects in each group. Segmentation maps from ACE were voxelized and warped to ABA 25-μm resolution. Panels show a randomly selected subject from the pargyline group. (B) Atlas plot of the average intensity difference of vCATCH labeling intensity of vehicle vs. pargyline-yne treated samples. The greatest labeling intensity elevation is denoted in yellow. (C) Atlas plot of the average segmented neuronal density (number of cells per mm^3^) of vehicle vs. pargyline-yne treated samples. The greatest density elevation is denoted in yellow. (D) Atlas plot of the average [-log(p-value)] of segmented neuronal density in vehicle vs. pargyline-yne treated samples. The greatest statistical significance is denoted in yellow. (E) MAO-A and - B immunostaining in pargyline-yne positive regions. (F) Histology validation of pargyline-yne binding in the LC. The boundary of LC and B is identified by NeuN and tyrosine hydroxylase (TH) staining using a published protocol^86^. Pargyline-yne binding is restricted in LC, suggesting that “B” was likely a false positive from ACE due to its small size. Scale bars: 200 μm (E and F).

**Figure S3:**
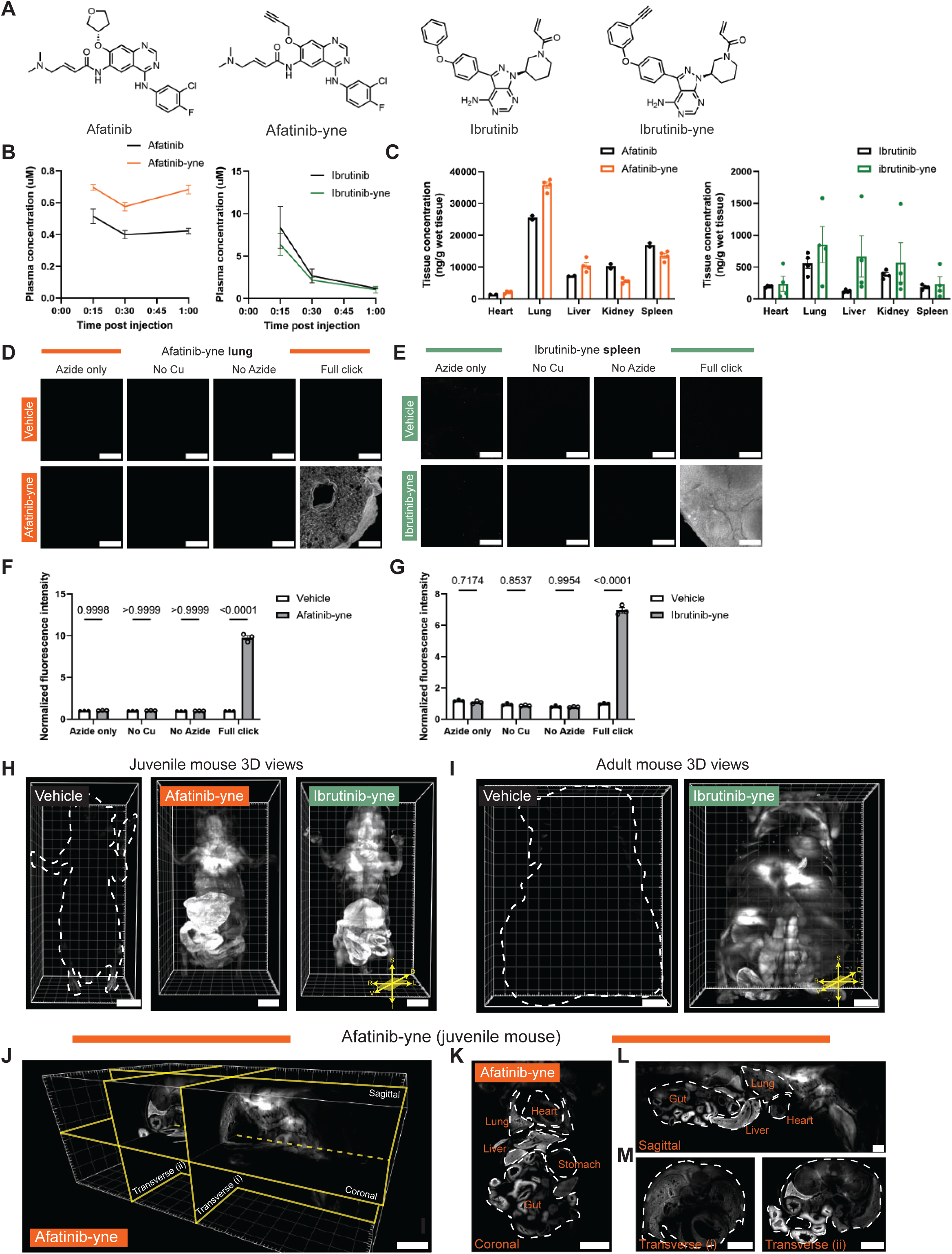
Whole body vCATCH visualization of TKI binding, related to Figure 3. (A) Chemical structures of afatinib, ibrutinib, and their alkyne analogs. (B) PK studies of plasma drug concentrations. Plasma samples taken 15, 30 and 60 min after drug injection (10 mg/kg). Two parental afatinib-60 min samples were excluded as outliers. n=4 mice for all time points except parental afatinib at 60 min (n = 2 mice). (C) PK studies of tissue drug concentrations at 60 min post injection (10 mg/kg). n=4 mice for afatinib-yne, ibrutinib, and ibrutinib-yne; n=2 mice for afatinib. (D-E) Click reaction specificity validation in afatinib-yne treated lung (D) and ibrutinib-yne treated spleen (E). (F-G) Quantifications of click labeling intensity in afatinib-yne-treated lung (F) and ibrutinib-yne treated spleen (G). Intensity normalized to vehicle full click condition. Three independent tissues were used for quantification. Two-way ANOVA with Šidák multiple comparisons test. (H) Representative 3D lightsheet image volume of whole-body drug distribution (10 mg/kg, 1h, i.p.) in juvenile vehicle, afatinib-yne, and ibrutinib-yne mice. D, dorsal; V, ventral; S, superior; I, inferior; R, right; L, left. (I) Adult whole torso 3D overview of mice treated with vehicle and ibrutinib-yne (10 mg/kg, 1h, i.p.). (J) Afatinib-yne distribution overviews with digital plane slicing in 3D volume image. (K-M) Coronal (K), sagittal (L), and transverse (M) views of mice injected with afatinib-yne as shown in Figure S3J. Data are plotted as mean ± SEM. Dashed lines indicate tissue boundary. Scale bars: 200 μm (D, E); 4000 μm (H, I, J, K); 2000 μm (L, M).

**Figure S4:**
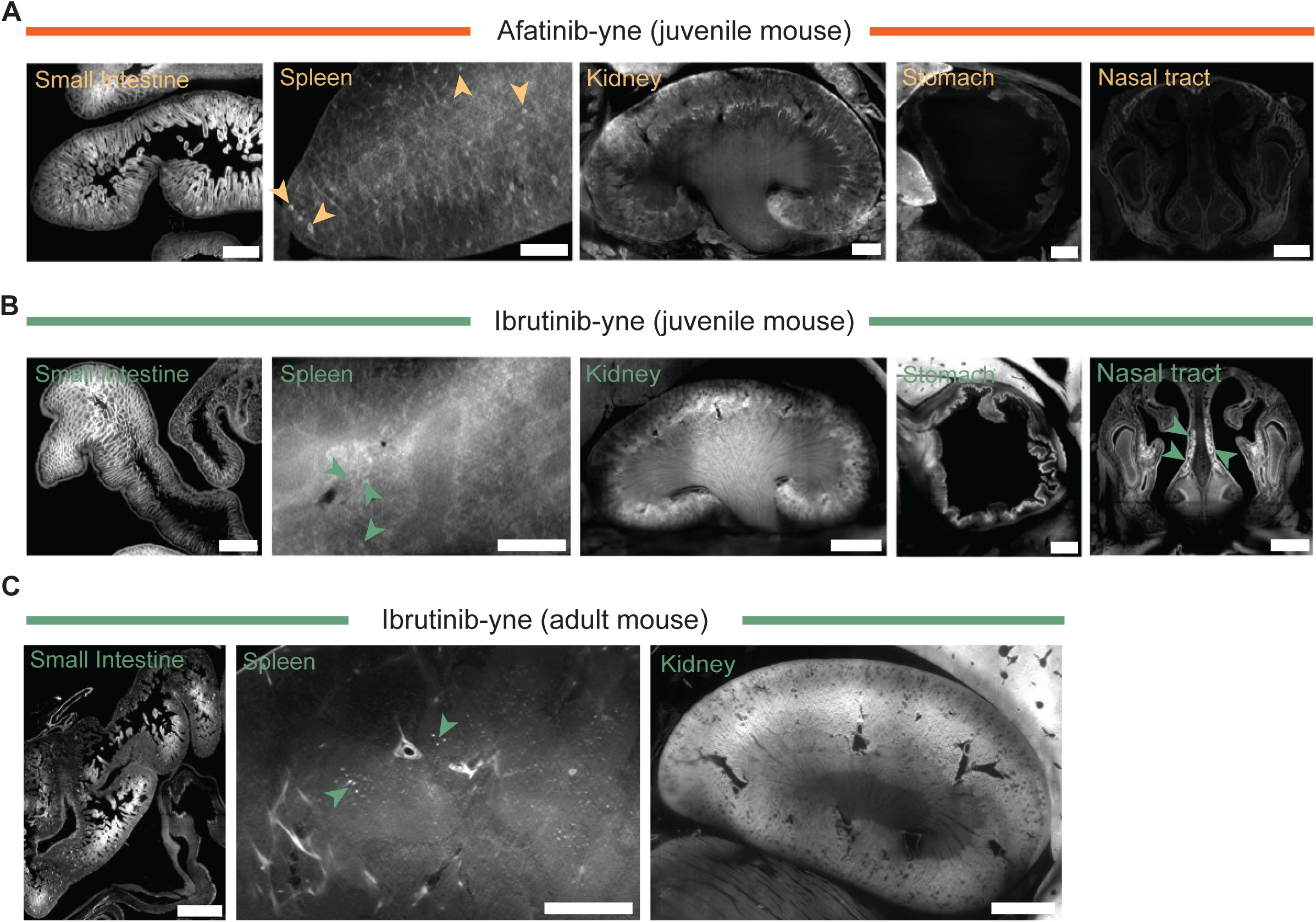
Organ level TKI binding, related to Figure 4. (A) Representative organ views from afatinib-yne treated mice. Afatinib-yne enrichment was observed in small intestine villi, spleen central arterioles (orange arrows), and kidney cortex. No significant drug enrichment observed in the stomach lining or nasal tract. Scale bars: small intestine, kidney, stomach, and nasal tract, 500 μm; spleen, 250 μm. (B) Representative organ views from ibrutinib-yne treated mice. Ibrutinib-yne enrichment was observed in small intestine villi, spleen central arterioles (green arrows), kidney, and inner stomach lining. Scale bars: kidney and stomach, 500 μm; small intestine and nasal tract, 300 μm; spleen, 200 μm. (C) Representative organ views from ibrutinib-yne treated adult mice. Ibrutinib-yne enrichment was observed in small intestine villi, spleen central arterioles (green arrows), and kidney. Scale bars: small intestine, 1000 μm; spleen, 500 μm; kidney, 1500 μm kidney.

**Figure S5:**
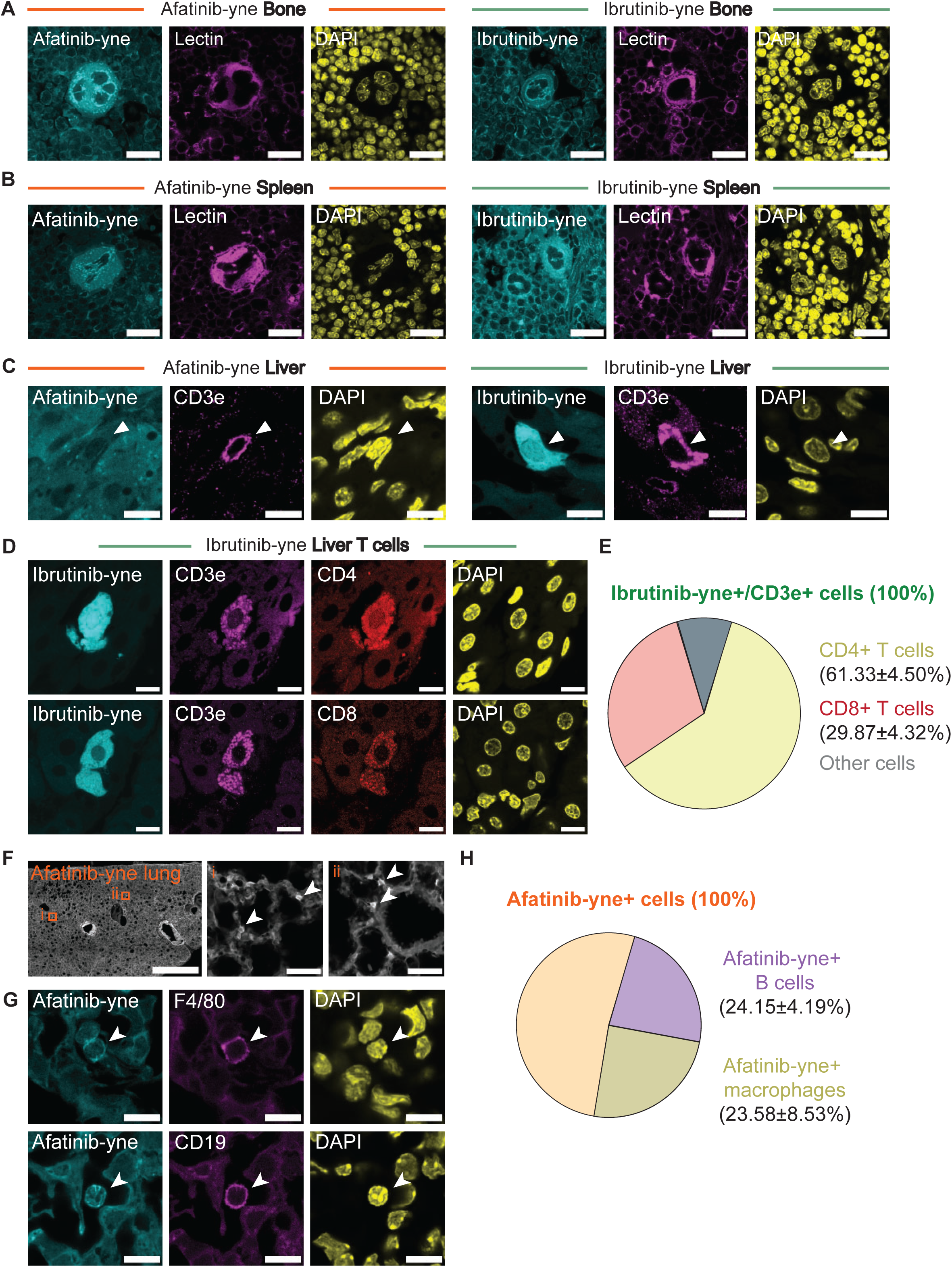
TKI positive cell type characterization, related to Figure 5. (A-B) Lectin staining in bone (A) and spleen (B) sections after CATCH labeling. Representative images showing lectin overlaps TKI enriched labeling structures. (C) High resolution afatinib-yne and ibrutinib T cell characterization in the liver. Representative images show CD3e+ cells that are afatinib-(left) and ibrutinib+ (right). Arrows indicate CD3e+ cells. (D) Cell type characterizations of ibrutinib-yne positive liver T cells. Samples are co-stained for helper T cells (CD4) and cytotoxic T cells (CD8). (E) Quantifications ibrutinib-yne positive T cell subtypes in the liver. n=4 biological replicates, 11-17 FOV were acquired for each mouse. (F) Global characterization of afatinib-yne binding in the lung. Zoom-in views showing discrete afatinib-yne enriched structures scattered throughout the lung. (G) High resolution afatinib-yne cell type characterization in the lung. Representative images showing afatinib-yne enriched F4/80 positive macrophages (top) and CD19 positive B cells (bottom). (H) Cell type characterization of afatinib-yne enriched cells in the lung. n=4 biological replicates, 5-6 FOV were acquired for each mouse. All values are shown as mean ± SEM. Detailed cell type quantification is in Supplementary Table 1. Scale bars: 20 μm (A and B); 10 μm (C, D, and G); 1000 μm (F, global view); 50 μm (F, zoomed-in view).

## STAR METHODS

Key Resources Table

## EXPERIMENTAL MODEL AND SUBJECT DETAILS

### Mouse model

Mice were group-housed on a 12-hr light-dark cycle and fed a standard rodent chow diet. Both male and female mice were used for the study. Six to nine-week-old WT C57BL6J mice were used for vCATCH method development in 100-micron sections and hemispheres. One-week-old and eight-week-old WT C57BL6J mice were used for vCATCH in the whole mouse body. All experimental protocols were approved by the Scripps Research Institute Institutional Animal Care and Use Committee and were in accordance with the guidelines from the NIH.

### Sample collection

Alkyne drugs were administered to mice in a vehicle of 10% DMSO and 2% Tween-80 in saline for i.p. injections at the indicated dose. 1 hour (for most direct labeling experiments) or 4 hour (for pretreatment test) after injections, mice were heavily anesthetized with isoflurane and then transcardially perfused with ice-cold PBS, followed by ice-cold 4% PFA solution. Mouse organs were carefully dissected and then post-fixed in 4% PFA solution overnight at 4℃. Tissues were washed with PBS. Mouse brains were embedded in 2% agarose and sectioned with a vibratome (Leica VT1000S) at 100-micron thickness. For hemispheres, brains were cut along the midline with a razor knife. Peripheral organs from one-week-old mice were dehydrated in 30% sucrose at 4℃ prior to sectioning. Samples were then carefully dried with a kimwipe and embedded in O.C.T. compound. After reaching equilibrium at -20℃, samples were cut at 100-micron thickness with a cryostat (Leica CM 1860). All obtained samples were stored in PBS with 0.02% sodium azide at 4℃ for further processing.

For mice used in whole-body drug distribution labeling, the skin was carefully removed immediately after perfusion, and the peritoneum was cut down the midline to facilitate clearing and click reactions. Mice were post-fixed in 4% PFA solution for two days at 4℃ followed by 7-day decalcification treatment in 10% EDTA/15% Imidazole at 4℃. Then, samples underwent decoloring treatment with 25% N, N, N’, N’-Tetrakis(2 Hydroxypropyl)ethylenediamine (Quadrol) in 1x PBS at 37°C with gentle agitation for 7 days. After PBS washes to remove Quadrol, mouse whole bodies were stored in PBS with 0.02% sodium azide at 4℃ for further processing.

For mice used in PK studies, after anesthesia, blood was collected via cardiac puncture and immediately stored in a heparin tube and placed on ice. Plasma was separated by centrifugation at 5,000 rpm for 5 min at 4°C. Next, 150 μL of a 2:1 mixture of acetonitrile/methanol was added to 50 μL of plasma. The mixture was then centrifuged at 15,000 rpm for 10 min at 4°C, and the supernatant was transferred to a liquid chromatography-mass spectrometry (LC-MS) vial. Tissue samples were dissected out and weighed. Next, tissues were homogenized in water at 250 mg/ml using a Homogenizer at 4°C. The mixture was then centrifuged at 15,000 rpm for 10 min at 4°C. 50 μL of the supernatant was then collected, and 150 μL of 2:1 mixture of acetonitrile/methanol was added to the supernatant. The mixture was then centrifuged at 4°C at 15,000 rpm for 10 min, and the supernatant was transferred to an LC-MS vial. To generate a standard curve, compounds were dissolved in a 2:1:1 mixture of acetonitrile/methanol/water at 10, 1, 0.1, 0.01, 0.001 μM for MS analysis.

### Tissue clearing

#### CLARITY

100-micron sections were cleared with CLARITY as previously described^22,87^. Briefly, samples were incubated in A1P4 hydrogel (1% acrylamide, 0.125% Bis, 4% PFA, 0.025% VA-044 initiator (w/v), in 1X PBS at 4℃ for CLARITY embedding. After overnight incubation with gentle agitation, samples were then degassed and polymerized at 37℃ for 4 hours with gentle agitation. Samples were removed from hydrogel and washed with 6% PBS-SDS (pH=7.0) at 37℃ for two days. After clearing, samples were washed with PBST (pH=7.0 with 0.2% Triton-X100, same for the following) three times (10 min each time) at RT to remove residue SDS. Samples were briefly washed with PBS and then stored in PBS with 0.02% sodium azide at 4℃.

#### HYBRiD

Mouse hemispheres and bodies were cleared with HYBRiD as previously described^27^. In brief, samples were dehydrated in an increasing THF/25% Quadrol gradient (50%, 70%, 80%, 100%, 100%; 1 hour per step for hemispheres, 3-12 hours per step for whole bodies), delipidated three times in DCM (1 hour each for hemispheres, 3-12 hours each for whole bodies), and rehydrated in a decreasing THF/25% Quadrol gradient (100%, 100%, 80%, 70%, 50%; 0.5 hour per step for hemispheres, 1.5-6 hours per step for whole bodies). After clearing, samples were extensively washed with PBS to remove residual organic solvent before hydrogel embedding. Samples were equilibrated in A1P4 hydrogel (3 days for hemispheres, 7-10 days with one refreshment for whole bodies) at 4℃ on a shaker, then degassed and polymerized at 37°C for 4 hours with gentle agitation. Thereafter, samples were passively cleared with LiOH Boric Buffer with 6% SDS (pH=9) until transparent (1 week for hemispheres, 6-10 weeks for whole bodies). Samples were extensively washed in PBST to remove residual SDS and then stored in PBS with 0.02% sodium azide at 4℃.

### Click labeling in tissues

#### CATCH in 100-micron sections

100-micron sections were labeled with an updated CATCH protocol. Unless otherwise noted, sections were incubated in 10 mM CuSO_4_ in H_2_O at RT overnight. Sections were then transferred to CATCH reaction buffer containing 5 μM AZDye 647-picolyl azide, 150 μM CuSO4, 300 μM BTTP, 2.5 mM NaAsb, and 10% DMSO in 1X PBS. After 1 hour of reaction in the dark with minor agitation, samples were washed with buffer containing 4 mM EDTA in PBST (PBST-EDTA, same for the following) at RT for 3x10 min. Tissue samples were then stained with DAPI (1:3000 dilution in PBST from 10 μM stock) for 15 mins at RT. Samples were then either used for refractive index (RI) matching by RapiClear and confocal imaging or proceeded to further immunolabeling for cell-type registration.

#### vCATCH in hemispheres

For hemisphere labeling, each hemisphere was first pre-incubated with 25 mL of 10 mM CuSO4 in H_2_O for one day at RT for PRCS. Hemispheres were transferred to vCATCH incubation buffer (5 μM AZDye 647-picolyl azide, 1 mM CuSO4, 2 mM BTTP, 10% DMSO in 1X PBS, 3 mL for each hemisphere) for 1 day at RT. Hemispheres were then transferred to the newly prepared full vCATCH reaction buffer (5 μM AZDye 647-picolyl azide, 1 mM CuSO4, 2 mM BTTP, 25 mM NaAsb, 10% DMSO in 1X PBS, 3 mL for each hemisphere) for click labeling. The vCATCH reaction buffer was refreshed every hour for two rounds of RIR. Hemispheres were washed with PBST-EDTA at RT. The washing buffer was refreshed at the end of the day and then daily until it became colorless. The hemispheres were then washed with PBST for 1 hour at RT. After washing, hemispheres were cut into 100-micron coronal sections for labeling depth characterizations. Alternatively, hemispheres were carefully dried with a Kimwipe and then incubated in 3 mL EasyIndex for RI matching and lightsheet imaging.

#### vCATCH in the whole mouse

Cleared mouse bodies were placed in 500 mL per mouse of 10 mM CuSO_4_ in H_2_O with agitation for 4-7 days at RT with daily buffer refreshes. After PRCS, whole-body samples were individually incubated in 10-40 mL of vCATCH incubation buffer for 3-5 days on a shaker at RT with daily buffer changes. Samples were then transferred to vCATCH reaction buffer (10-40 mL each) to initiate the reaction. Five to eight successive rounds of 1-hour click reactions were performed to thoroughly label the entire tissue depth. After click labeling, samples were washed with PBST-EDTA with daily refreshments until the buffer was colorless. After an additional day of PBST wash, mouse bodies were carefully dried with a Kimwipe and then thoroughly immersed in 25-40 mL EasyIndex at 37°C for thorough RI matching in preparation for lightsheet imaging.

### Immunostaining

CATCH-labeled 100-micron sections were incubated with primary antibodies 1:200 diluted in PBST overnight at 4℃. Samples were then washed with PBST 3 x 30 min at RT prior to incubation with secondary antibodies diluted 1:600 in PBST. Samples were then incubated with secondary antibodies 1:600 diluted in PBST. Samples were incubated with secondary antibodies overnight and then washed with PBST 3 x 30 min at RT.

### Confocal microscopy

100-micron samples were mounted with RapiClear for RI matching and imaged with an Olympus FV3000 confocal microscope. To characterize CATCH labeling depth in 100-micron sections, samples were imaged at the primary somatosensory cortex layer V with a 10X, 0.6 NA, water immersion objective (XLUMPlanFI, Olympus) at 0.414 micron/pixel resolution and a step size of 10 microns. To examine hemisphere labeling depth and global drug distribution within individual organs, samples were imaged with a 10X, 0.6 NA, water immersion objective (XLUMPlanFI, Olympus) at 2.49 microns/pixel resolution and a step size of 10 microns. For cell type characterizations, samples were imaged at the top surface with a 40X, 1.25 NA, silicone oil objective (UPlanSApo, Olympus) at 0.094 micron/pixel resolution and a step size of 3 microns. All confocal data was saved in TIFF format for further analysis.

### Lightsheet microscopy

Whole hemisphere/body samples were imaged using a lightsheet microscope from LifeCanvas Technologies. RI-matched whole hemisphere and body samples were mounted in 1% agarose/Easy Index for imaging. Lightsheet imaging was performed in a SmartSPIM chamber with Easy Index matched immersion oil (RI=1.52). A 3.6x objective was used with 0.28 NA, 1.8μm/1.8μm/2μm *xyz* voxel size. Imaging was performed by illuminating samples along the bilateral plane of symmetry. Lightsheet raw data was then destriped, stitched, and saved as TIFF format for further conversion.

### Image processing and analysis

#### Hemisphere labeling depth characterization

Hemispheres were sectioned coronally as 100 micron sections and imaged to characterize vCATCH labeling depth. Images were first processed as maximum intensity projection. A rectangular area (805 x 50 pixels, 2001.17 x 136.73 microns) was drawn from the cortical outer layer inwards (from tissue surface to tissue center) as indicated in Figure 1A. The signal profile was plotted using the ‘Plot Profile’ function in ImageJ as reported^27,88,89^. The fluorescence signal profile was then normalized by the intensity value at the start of the line.

#### Thin section labeling depth characterization

CATCH labeling intensity across the 100-micron section was quantified as previously reported^21,27,88^. Briefly, a 150 x 150 pixel (62.1 x 62.1 micron) ROI surrounding a labeled neuron was generated at z = 10 μm and 50 μm at cortex layer V, and an auto threshold was applied to measure the mean drug-positive pixel intensity as *I*_signal_. The mean intensity of the remaining pixels was used *I*_background_. The mean labeling intensity was defined as *I*_labeling_=*I*_signal_-*I*_background_.

#### Cell type-specific characterization

CD3e, F4/80, CD4, CD8, and CD19 positive cells were characterized by immune staining. Hepatocytes were identified based on their large size (20-30 microns), round/oval-shaped nucleus, and hexagonal morphology by well-trained experts. To minimize bias, individual fields of view (FOV) were first acquired based on the DAPI and the immunostaining channel without knowledge of the CATCH/drug channel (with the exception of CD3e+ibrutinib+ cells, where the CD4 and CD8 channel were kept blind during imaging acquisition) to assign cell types. The CATCH positivity was post-hoc assigned to individual cells after their cell types have been determined, followed by overlapping quantifications. For intensity characterizations, individual cells were selected along the boundary and cropped out for quantification. A threshold was applied in the drug channel to analyze CATCH labeling intensity in individual cells of interest. The full dataset of each cell population can be found in Supplementary Table 1.

#### Global drug intensity characterization

To measure global drug intensity across intact thin tissue sections, confocal images were first processed as maximum intensity projections. A threshold was then applied in the counterstain channel (DAPI or target immunostaining) to cover the whole tissue or target positive pixels. The threshold area was selected as tissue containing pixels. The average drug channel intensity in the selected pixels was then quantified as global drug intensity.

#### Volumetric drug binding visualization

Lightsheet imaged hemispheres and whole body images were converted into Imaris files and visualized via Imaris 10.2.0. The hemisphere drug labeling video was generated by Imaris. For VR-based whole-body drug visualization, data was converted by syGlass, and videos were made with a Meta Quest 3 glass. For hemisphere coronal and transverse views, TIFF files were imported as image sequences and then resliced by Fiji ImageJ. For drug distribution visualization across organs, TIFF files were imported as image sequences and sampled at 100-micron step size. Individual organ of interest was then cropped out for visualization.

#### Automated Brain-wide Cell Mapping & Signal Intensity Calculation

We employed an AI-based Cartography of Ensembles (ACE) pipeline to generate whole-brain segmentation maps of pargyline-yne+ cells^53^. ACE is an open-source 3D deep learning pipeline (available through the MIRACL platform^90^; https://miracl.readthedocs.io/) to map local cell activations using tera-voxel size light sheet microscopy datasets. Given the significant differences in morphological features between pargyline-yne+ cells and cFos+ cells (which the ACE models were originally trained on), we first fine-tuned the ACE vision transformer (ViT) model. For fine-tuning, we randomly selected 225 image patches of 963 voxels each (0.17x0.17x0.19 mm^3^) from two subjects in the dataset and generated ground truth using the Labkit annotator^91^ in ImageJ/Fiji software^92^. We fine-tuned the ViT model on half of these patches and used the other half for evaluation. The model was fine-tuned for 200 epochs using the Adam optimizer^93^ with an initial learning rate of 0.0001 and an equally weighted Dice-Cross Entropy loss function. To account for artifacts in the dataset, the cross-entropy calculation for “background” voxels was weighted 2× higher than for voxels annotated as pargyline-yne+ cells. The best model, which achieved an average Dice score of 0.7 on the validation dataset, was used to generate segmentation maps for the entire dataset. Subsequently, a shape filter was applied to the segmentation maps to eliminate false positives such as vasculature traces in cortical regions using the shape filter plugin in ImageJ/Fiji^94^. To distinguish individual cells, a connected component analysis was applied to the 3D segmentation maps of each subject. Following segmentation, we automatically registered the light sheet hemisphere data to the ABA using the MIRACL platform^54,90^. The accuracy of the registration and warping were evaluated using quality control checkpoints. Deformation fields obtained from the registration algorithm were used to warp the ABA labels to the native space of each subject. Finally, we computed the number of cells within each region at depth 6 (labels grouped to a maximum atlas ontology depth of 6, by combining finer labels under their parent labels), by identifying the coordinates of the center of each cell.

To map significant localized group-wise differences in pargyline-yne+ cells in an atlas-agnostic manner, we employed ACE’s cluster-wise threshold-free cluster enhancement permutation test using a group-wise ANOVA. The segmentation maps obtained by the deep learning models were downsampled to ABA 10-μm resolution while minimizing information loss by applying a convolution-based voxelization procedure to the whole-brain segmentation maps (based on our prior work^90^). Following voxelization, segmentation maps were aligned with the ABA at a 10-μm resolution using deformation fields obtained via the registration module. The voxelated and warped segmentation maps were then passed to the ACE statistical module (using default parameters), which produced a cluster-wise p-value map representing significant (alpha < 0.05 after correction) localized cell activation between the pargyline-yne treated and control groups.

We computed signal intensity information for each ABA label using the “*miracl lbls stats*” function in our MIRACL platform, which processes NIfTI images and their registered warped ABA labels for each subject. This function extracts key intensity metrics, such as the average, minimum, and maximum voxel intensity, across the input volumes for each label. We further summarized it by grouping labels at depth 6 using the atlas ontology.

#### Brain-wide drug binding heatmap visualization

Brain regions with positive cell counts in all 5 pargyline-yne-treated hemispheres were included for heatmap generation. Cell density (number of cells per mm^3^) was obtained by dividing cell count values by the volume of the brain region. Intensity and cell density were then normalized by the average values in individual hemispheres. Heatmaps were generated using the Python function ‘sns.heatmap’^95^, with each region displaying the normalized intensity and density across different subjects. The height of each subplot, corresponding to different regions, is proportional to the number of areas within that region. Code for generating heatmaps is available via https://github.com/yelabscripps/vCATCH.

### Drug PK analysis

Targeted measurements were performed using an Agilent 6470 triple quadrupole LC-MS instrument. MS analysis was performed using AJS in positive mode. The AJS source parameters were set as follows: the dry gas temperature was set at 250 °C with a gas flow of 12 l/min and the nebulizer pressure at 25 psi; the sheath gas temperature was set to 300 °C with the sheath gas flow set at 12 l/min; and the capillary voltage was set to 3,500 V. Separation of compound was conducted using a ZORBAX RR Eclipse Plus C18 95Å LC column (Agilent 959961-902) with reversed-phase chromatography and the column temperature was maintained at 30 °C. Mobile phases were as follows: buffer A, 100% water with 0.1% formic acid; buffer B, 100% acetonitrile with 0.1% formic acid. The flow rate for each run started at 95% A for 9 minutes at 0.7 ml/min, followed by a gradient starting at 95% A, changing linearly to 5% A / 95% B over the course of 12 minutes at 0.7 ml/min. The flow rate was maintained at 5% A / 95% for 6 minutes at 0.7 ml/min. The last 3 minutes consisted of a re-equilibration back to 95% A / 5% B at 0.7 ml/min. Multiple reaction monitoring was performed for the indicated chemicals with the listed dwell times, fragmentor voltage, collision energies, cell accelerator voltages and polarity. We selected previously reported transitions for the quantification of ibrutinib^96^ and afatinib^97^. The MS ionization parameters for the targeted metabolomics are presented in Supplemental Table 1. Quantification of the compound concentrations was performed by generating an external standard curve with known concentrations of each compound. Compound standards were analyzed alongside the biological samples using the same targeted triple quadrupole LC/MS method in the same run. A calibration standard curve generated from the compound standard concentrations and total peak areas were used to calculate the concentrations of each compound. It is worth noting that, contrary to the high liver distribution from the autoradiography study^72^, our PK data showed overall lesser ibrutinib liver enrichment (Figure S3C). The caveat may be due to the different detection analytes (isotope tag vs. intact free drug). And the observed ibrutinib liver enrichment may reflect strong local metabolism and potential off-target binding.

## SUPPLEMENTAL INFORMATION

**Supplementary Table 1: Raw data summary and key MS parameters. Related to Figure 1C, 2D, 2E, 5I, 5J, 5L, 5M, 5O, 5P, S1C, S3B, S3C, S3F, S3G, S5E, S5H.**

**Supplementary Video 1: 3D rendering of pargyline-yne binding across the whole mouse hemisphere. Related to Figure 1D**.

**Supplementary Video 2: 3D rendering of afatinib-yne binding across the whole mouse body. Related to Figure S3H, S3J.**

**Supplementary Video 3: 3D rendering of ibrutinib-yne binding across the whole mouse body. Related to Figure S3H, 3E, S3H.**

